# Higher-order sensorimotor circuit of the brain’s global network supports human consciousness

**DOI:** 10.1101/2020.09.22.308072

**Authors:** Pengmin Qin, Xuehai Wu, Changwei Wu, Hang Wu, Jun Zhang, Zirui Huang, Xuchu Weng, Zengxin Qi, Weijun Tang, Tanikawa Hiromi, Jiaxing Tan, Sean Tanabe, Stuart Fogel, Anthony G. Hudetz, Yihong Yang, Emmanuel A Stamatakis, Ying Mao, Georg Northoff

**Affiliations:** Key Laboratory of Brain, Cognition and Education Sciences, Ministry of Education; School of Psychology, Center for Studies of Psychological Application, and Guangdong Key Laboratory of Mental Health and Cognitive Science, South China Normal University, Guangzhou, Guangdong, 510631, China; Pazhou Lab, Guangzhou, 510335, China; Department of Neurosurgery, Huashan Hospital, Shanghai Medical College, Fudan University, Shanghai, China; Neurosurgical Institute of Fudan University, Shanghai Clinical Medical Center of Neurosurgery, Shanghai Key laboratory of Brain Function Restoration and Neural Regeneration, Shanghai, China; State Key Laboratory of Medical Neurobiology and MOE Frontiers Center for Brain Science, School of Basic Medical Sciences and Institutes of Brain Science, Fudan University, Shanghai, China; Research Center for Brain and Consciousness, Taipei Medical University, Taipei, Taiwan; Graduate Institute of Humanities in Medicine, Taipei Medical University, Taipei, Taiwan; Shuang-Ho Hospital, Taipei Medical University, New Taipei, Taiwan; Department of Anesthesiology, Fudan University Shanghai Cancer center,Shanghai, China; Department of Anesthesiology and Center for Consciousness Science, University of Michigan, Ann Arbor, MI, USA; Institute for Brain Research and Rehabilitation, South China Normal University, Guangzhou, Guangdong, China; Radiology Department, Shanghai Huashan Hospital, Fudan University, Shanghai, China; School of Psychology, University of Ottawa, Ottawa, Canada; Neuroimaging Research Branch, National Institute on Drug Abuse, Intramural Research Programs, National Institutes of Health, Baltimore, USA; Division of Anaesthesia, School of Clinical Medicine, University of Cambridge, Cambridge, UK; Institute of Mental Health Research, University of Ottawa, Ottawa, Ontario, Canada; Mental Health Centre, Zhejiang University School of Medicine, Hangzhou, China

**Keywords:** Anesthesia, Degree centrality, Inferior parietal lobule, rapid eye movement sleep, higher-order sensorimotor circuit, disorders of consciousness

## Abstract

The neural correlates of consciousness, defined as the minimum neuronal mechanisms sufficient for any conscious percept, are usually subject to different interpretations depending on whether one uses measures of local or global brain activities. We argue that the local regions may support consciousness by serving as hubs within the brain’s global network. We adopt a unique functional magnetic resonance imaging resting state dataset that encompasses various conscious states, including non-rapid eye movement (NREM)-sleep, rapid eye movement (REM)-sleep, anesthesia, and brain injury patients. Using a graph-theoretical measure for detecting local hubs within the brain’s global network, we identify various higher-order sensory and motor regions as hubs with significantly reduced degree centrality during unconsciousness. Additionally, these regions form a sensorimotor circuit which correlates with levels of consciousness. Our findings suggest that integration of higher-order sensorimotor function may be a key mechanism of consciousness. This opens novel perspectives for therapeutic modulation of unconsciousness.

## Introduction

Consciousness is a core mental feature and the neural correlates of consciousness (NCC) (Crick & Koch, 2003) have been the focus of intensive investigation in neuroscience (Bachmann, 2015; Koch, Massimini, Boly, & Tononi, 2016; Tononi, Boly, Massimini, & Koch, 2016). Studies employing various methodologies including functional magnetic resonance imaging and multichannel electroencephalography so far have yielded conflicting views on whether resting state and task-evoked activity of specific sensory and cognitive brain regions such as the posterior “hot zone” (Koch et al., 2016) or the dorsolateral prefrontal cortex may serve as the NCC (Dehaene & Changeux, 2011). In addition to specific regions and regional specialization, the long-distance communication among remote brain regions as well as global activity changes have been emphasized (Huang et al., 2016; Luppi et al., 2019; Tanabe et al., 2020; Tang et al., 2017). Efforts have been directed to identifying the particular regions of the brain’s global functional network, in graph theoretical terms called “hubs” may be critical in supporting consciousness; hubs or nodes reflect local activity in specific regions that strongly modulate the brain’s global activity (Boly et al., 2017). While recent approaches identified the dynamic nature of the NCC (Demertzi et al., 2019), the regions and circuit that support consciousness within the brain’s global functional network remain yet unclear.

Insights into the NCC as a state have strongly benefited from investigations of abnormal behavioral and brain states characterized by reduced or absent consciousness, e.g., in anesthesia (Hashmi et al., 2017; Kertai, Whitlock, & Avidan, 2012; Monti et al., 2013; Moon, Lee, Blain-Moraes, & Mashour, 2015), sleep (Horovitz et al., 2009; Houldin, Fang, Ray, Owen, & Fogel, 2019; Hu, Cheng, Chiu, & Paller, 2020; Larson-Prior et al., 2009), unresponsive wakefulness syndrome (UWS) and minimally conscious state (MCS) (Engemann et al., 2018; Monti et al., 2010; Schiff, 2015). One particularly powerful approach is to identify brain regions and networks whose activity patterns track with variations in the behaviorally defined states of consciousness.

Prior investigations, especially in UWS, have demonstrated abnormal activity in higher-order sensory and motor regions like supplementary motor area (SMA) (Owen et al., 2006) and inferior parietal lobule (IPL) (Zhang et al., 2018), while the stimulus evoked activity of primary sensory regions is largely preserved (Di et al., 2007). These studies suggest that sensorimotor integration is required for normal, healthy consciousness (O’Regan & Noe, 2001). The findings suggest a key role for a higher-order sensorimotor integration circuit in supporting consciousness. However, additional higher-order regions and functional networks like the salience network (Qin et al., 2015), default-mode network (Vanhaudenhuyse et al., 2010), and frontal-executive network (Stender et al., 2014) have also been implicated in states of reduced or absent consciousness. Some studies even suggest that the brain’s global spatial activity pattern that operates across the whole brain may be central in supporting consciousness (Tanabe et al., 2020). Taken together findings on local and global activity changes, we are confronted with the question how local and global activity can be reconciled in their support for consciousness – that is the main focus of our study. Specifically, we asked whether consciousness is supported by the existence of a higher-order sensorimotor integration circuit within the brain’s global functional network.

To address the above question, degree centrality (DC), a graph theoretical measure, is a useful tool. Based on the resting-state fMRI data, degree centrality allows measuring the relative importance and contributions of specific regions serving as nodes within the global overall network architecture by calculating the number of connections made to a node (voxel) (van den Heuvel & Sporns, 2013). This approach is suitable for calculating the functional integration (centrality) of hubs in a complex network such as the brain. Brain regions with high degree centrality and their functional connections are thought to allow for higher-order integration of different inputs like sensory and motor, i.e., higher-order sensorimotor integration (Bullmore & Sporns, 2012).

In this study, we, using resting-state fMRI, aim to identify the higher-order integration nodes forming a circuit within the brain’s global functional network. Importantly, we aim to show how such circuit can track behaviorally defined states of consciousness from normal wakefulness to pharmacological and neuropathological states where consciousness was presumed to be reduced or lost. We take advantage of a large dataset that included subjects in anesthesia, sleep, and patients with disorders of consciousness (DOC, including UWS and MCS). Most importantly, we also include a unique fMRI dataset on REM-sleep, where sensory input is maximally reduced and the ability of intentional movement generation/execution is lost while the subjects still maintain rich conscious experiences.

Investigating REM-sleep will thus provide a unique window into understanding the NCC by itself independent of sensory and motor processing. This is even more important given that REM-sleep will be helpful to exclude the possibility of effects caused by a lack of mobility, since unconscious states are also characterized by immobility (e.g. sleep, anesthesia, and UWS). REM-sleep on the other hand, also helps excluding the possibility of effects caused by a lack of environmental sensory inputs, since REM-sleep and unconscious states share the same disconnection from the external world.

Specifically, the present study is divided into two parts. The first part is exploratory and involves a sleep group without REM (including awake state, N1-sleep, and N3-sleep), an anesthesia group (including conscious wake and anesthesia states), and DOC patients (including UWS, MCS and a fully conscious patient group with brain lesions (BL)) (See Supplementary Table 1). All DOC patients were structurally preserved brain injury patients (Supplementary fig. 1 for structural brain images). For sleep group, N3-sleep was regarded as an unconscious state (deep sleep stage) (Laureys, 2005). The second part of our study focuses on validation of the regions obtained in the first part by involving a REM-sleep group (including awake, N3-sleep and REM-sleep). For the data-analysis, during the first exploratory part, the brain regions with significant degree centrality reduction during unconsciousness were discovered, and then further confirmed with the REM-sleep dataset. The functional connections between each of these brain regions were investigated in all levels ranging from consciousness to unconsciousness; that served the purpose to extract the functionally relevant circuit among different regions within the global functional network. Please see Fig.1 for the main idea of the manuscript and the Schemata of data processing.

**Fig. 1.**
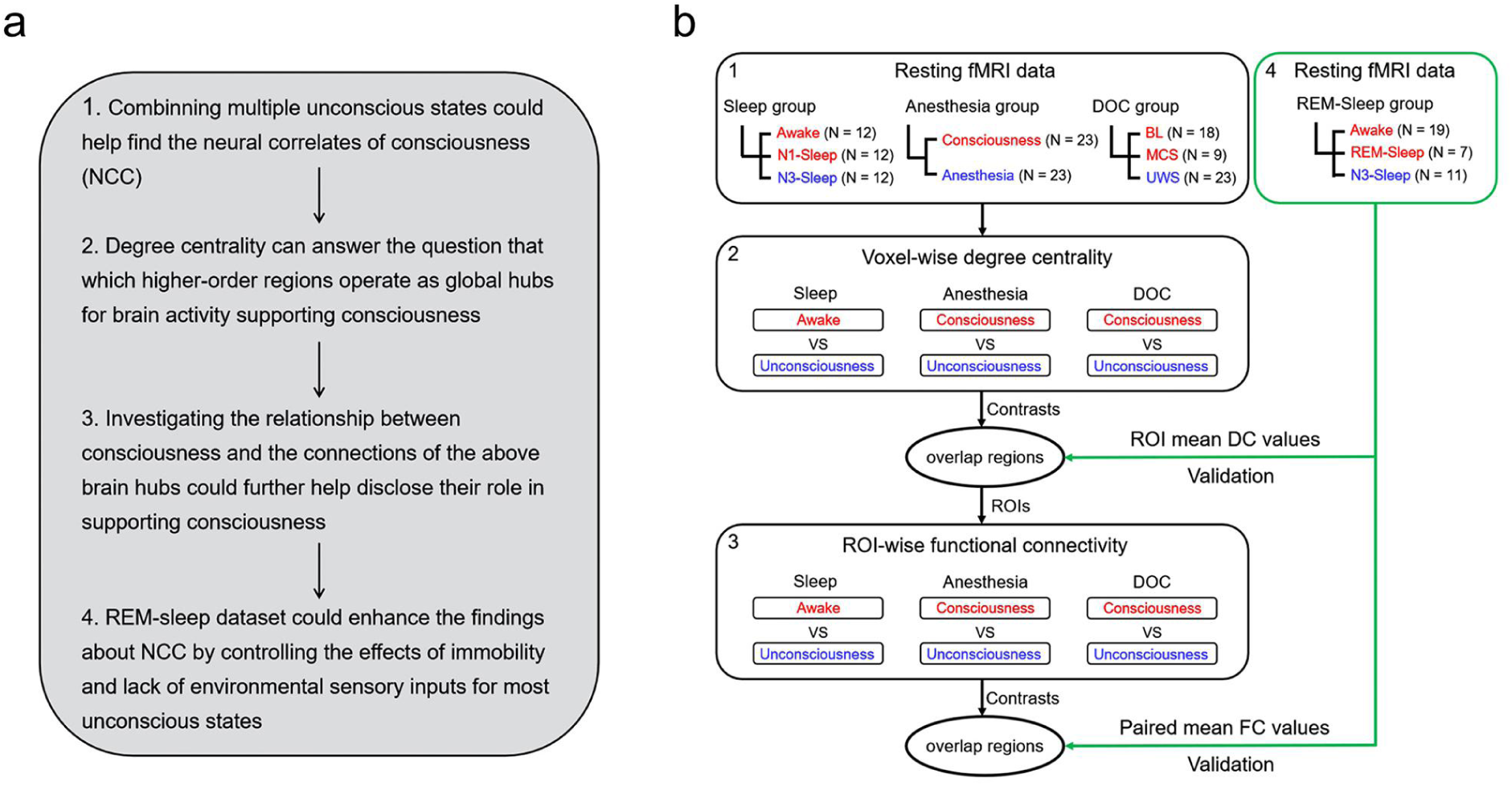
The schemata for experiment design and data processing. **a** The strategy of the experiment design and data analysis. **b** The schemata for experiment design. Red represents consciousness or reduced consciousness states; blue represents unconsciousness. The black square represents the exploratory datasets which were used to define the ROIs with degree centrality and functional connectivity changes during unconsciousness. The green square represents the validation datasets (REM-sleep group). DC values = degree centrality values, FC values = functional connectivity values. N means the sample size of each group. The digital number indicated the corresponding idea and data processing in panel **a** and **b**.

We identify various higher-order sensory and motor regions like bilateral supramarginal gyrus (SMG), supplementary motor area (SMA), supragenual/dorsal anterior cingulate cortex (SACC), and left middle temporal gyrus (LMTG) as important hubs within the brain’s global functional network. Additional analyses show that the same regions exhibit high degrees of functional connectivity among each other forming a higher-order sensorimotor circuit. Importantly, such sensorimotor circuit is observed across the different states entailing that it supports consciousness independent of the specific causes underlying the different states. Together, we demonstrate a higher-order sensorimotor circuit within the brain’s global network to support consciousness.

## Results

### Reduced degree centrality in unconsciousness

As a first step, whole brain voxel-wise degree centrality analysis was performed to build a degree centrality map (r > 0.3 was taken as one edge) (Huang et al., 2016). In order to reveal the degree centrality difference between complete absence of consciousness and presence (including both incomplete and complete) of consciousness, two contrasts were performed within the sleep group: N3-sleep vs. awake and N3-sleep vs. N1-sleep. One contrast was performed for the anesthesia group: anesthesia vs. consciousness. Two contrasts were performed for the DOC group: UWS vs. BL and UWS vs. MCS. The above contrasts showed significant degree centrality reduction during unconscious states (p < 0.05 FWE correction), which overlapped in SMA, left supramarginal gyrus (LSMG), right supramarginal gyrus (RSMG), supragenual anterior cingulate cortex (SACC) and left middle temporal gyrus (LMTG) (Fig. 2 and supplementary Table 2). These five regions did not show any degree centrality difference between N1-sleep and awake-states in the sleep group. LSMG showed significant differences between BL and MCS, but not other four regions (Supplementary fig. 2).

**Fig. 2.**
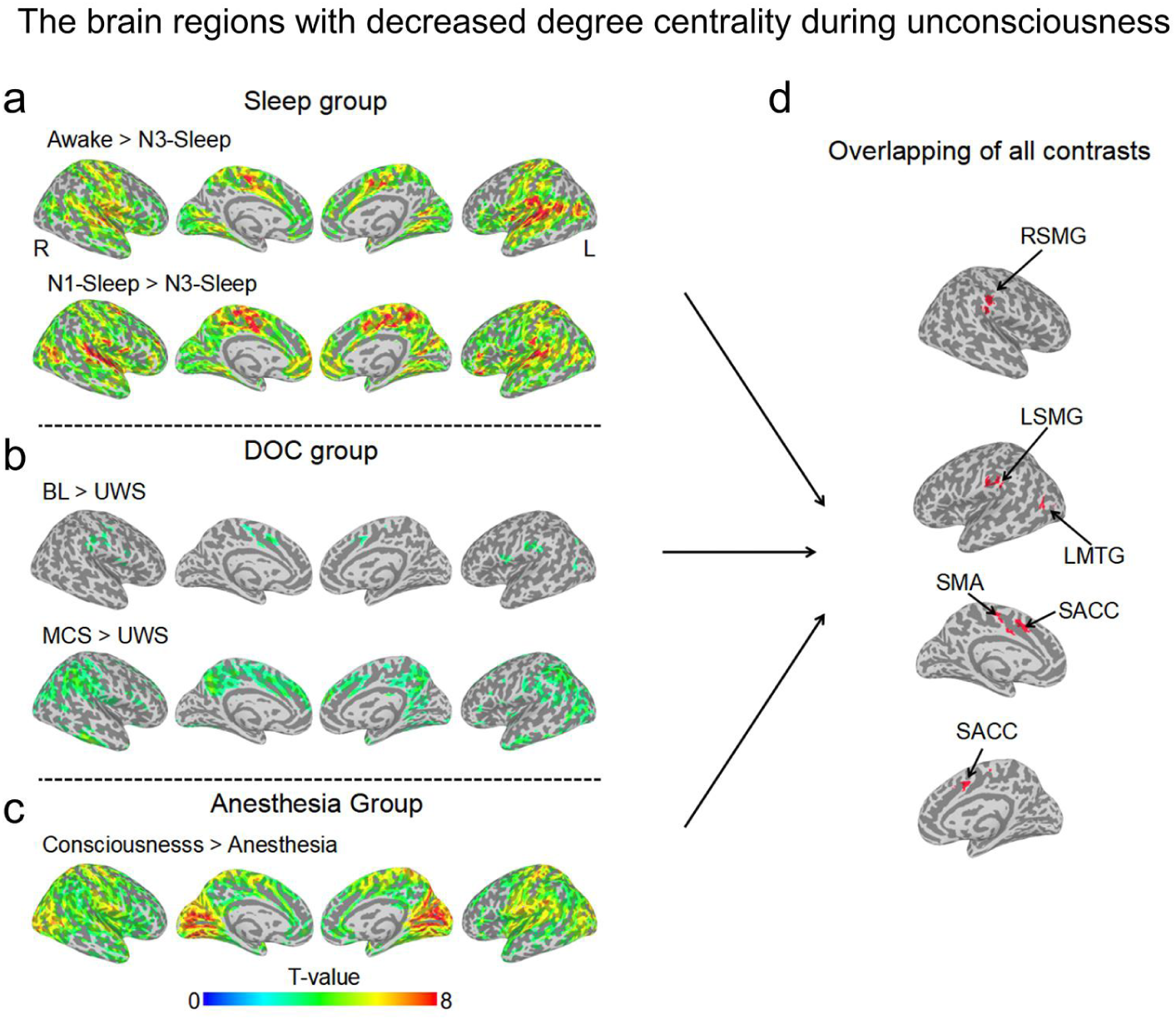
The brain regions with decreased degree centrality during unconsciousness. All the degree centrality contrasts for unconscious < conscious/reduced conscious states, (**a**) Sleep group, (**b**) DOC patients group and (**c**) anesthesia group. The activated clusters with (p < 0.005 uncorrected, volume > 20 voxels) are displayed. (**d**) The overlapped brain regions (volume > 20 voxels) of all the contrasts from panel **a**, **b**, and **c**. SMA = supplementary motor area; LSMG = left supramarginal gyrus; RSMG = right supramarginal gyrus; LMTG = left middle temporal gyrus; SACC = supragenual anterior cingulate cortex.

### Intact degree centrality in REM-sleep

The above five regions identified (SMA, SACC, bilateral SMG, and LMTG) were used as ROIs in REM-sleep datasets to test whether their degree centrality would be intact during REM-sleep. SACC showed a significant difference between states (H(2) = 14.98, *η*^2^ = 0.41, p < 0.001; post-hoc test showed N3-sleep < REM-sleep, p < 0.001, and N3-sleep < awake, p = 0.014). SMA showed a significant difference between states (H(2) = 14.01, *η*^2^ = 0.36, p < 0.001; post-hoc test showed N3-sleep < REM-sleep, p = 0.001, and N3-sleep < awake, p = 0.003). LSMG showed a significant difference between REM-sleep and N3-sleep states (H(2) = 6.94, *η*^2^ = 0.15, p = 0.031; post-hoc test showed N3-sleep < REM-sleep, p = 0.014, and N3-sleep < awake, p = 0.085). RSMG showed a significant difference between REM-sleep and N3-sleep states (H(2) = 8.53, *η*^2^ = 0.20, p = 0.014; post-hoc test showed N3-sleep < REM-sleep, p = 0.007, and N3-sleep < awake, p = 0.089). LMTG showed a significant difference between states (H(2) = 17.15, *η*^2^ = 0.46, p < 0.001; post-hoc test showed N3-sleep < REM-sleep, p < 0.001, and N3-sleep < awake, p < 0.001). All p values above were FDR corrected (Fig. 3).

**Fig. 3.**
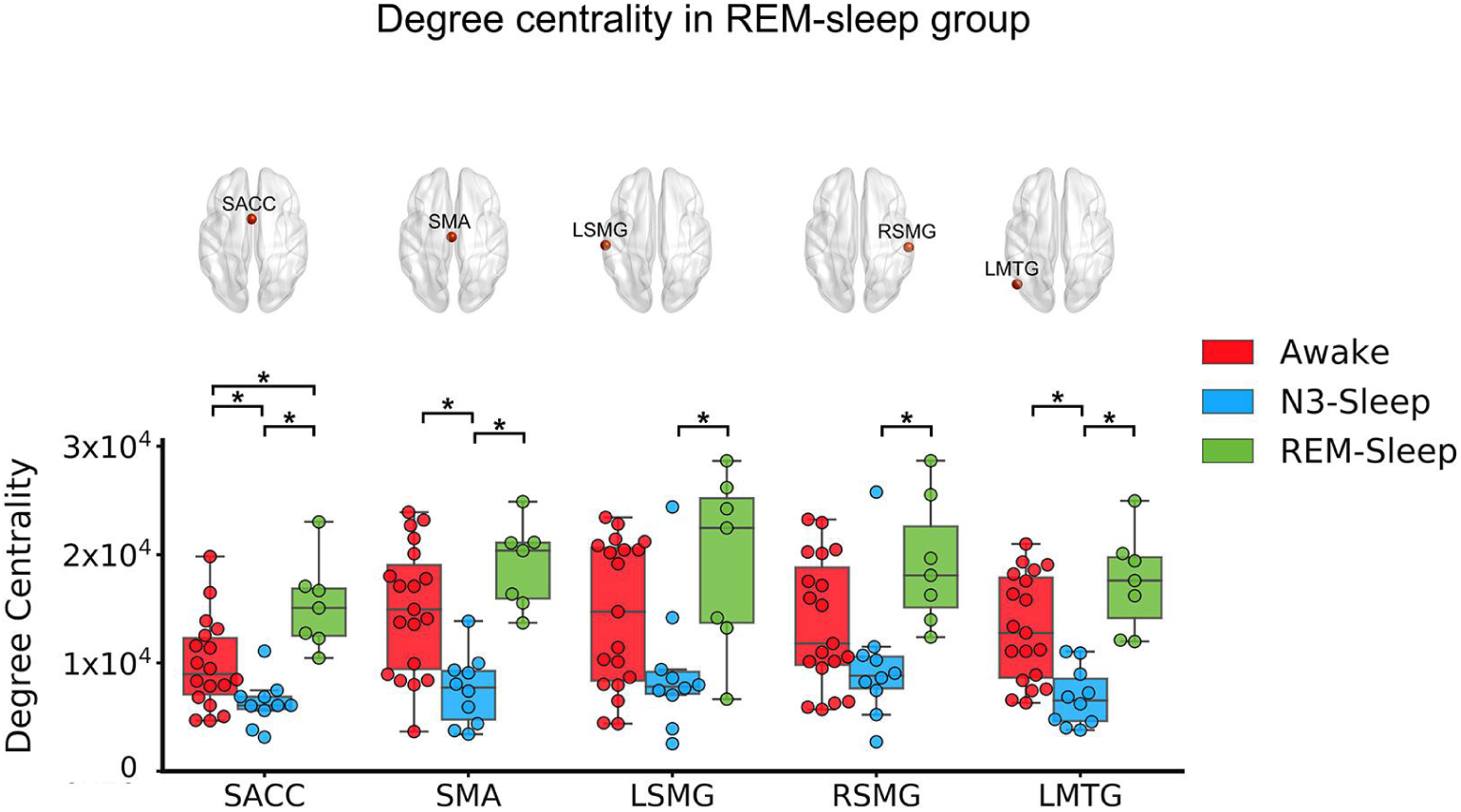
Degree centrality in REM-sleep group. The degree centrality (DC) values for awake, REM-sleep and N3-sleep states in SMA, SACC, LSMG, RSMG, and LMTG. * means p < 0.05 FDR corrected.

We then validated the above findings by performing degree centrality analysis with different edge thresholds (r > 0.2 and r > 0.4) (Dai et al., 2015). Our results showed that all findings remained consistent when using different thresholds (Supplementary fig. 3).

### ROI-based functional connectivity during consciousness

The degree centrality reflects the connectivity of one region with the whole brain. It is a quantitative measure but does not provide any regional connectivity information. To reveal the connectivity of each pair of regions among the five brain regions (ROIs obtained in the degree centrality analysis), as well as which connectivity showed reduction during unconsciousness, ROI-based functional connectivity was performed for all the states in each group: sleep group, anesthesia group, DOC group, and REM-sleep group. The results showed that all five regions had significant functional connectivity (z > 0.3095) during consciousness or awake-states within each group, except the functional connectivity of LMTG with SMA and SACC in anesthesia group. More interestingly, the strengths of the functional connectivity showed significant correlations between each pair of groups during consciousness or awake- states (Fig.4). These results indicated that the functional connectivity between each pair of regions, as well as its relationship with other functional connectivity, was stable during consciousness. Please see the functional connectivity values of the five regions in reduced consciousness and unconsciousness in Supplementary fig. 4.

**Fig. 4.**
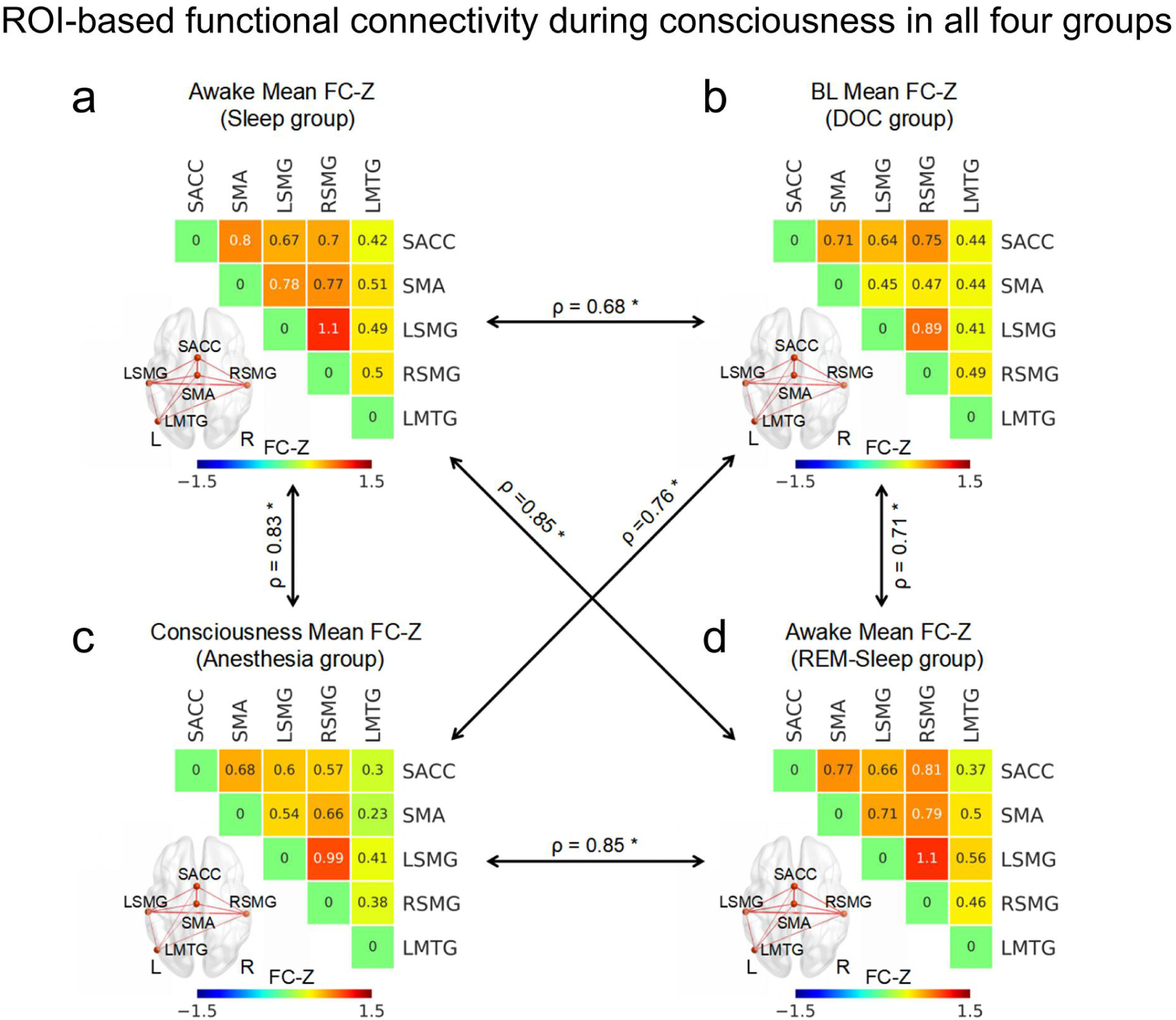
ROI-based functional connectivity during consciousness in all four groups. The ROI- based functional connectivity z-values for the awake-state in the sleep group (**a**), for BL patients in the DOC group (**b**), for the conscious state in the anesthesia group (**c**), and for awake-states in the REM-sleep group (**d**). The black arrow represented the relationship of these functional connectivity between each pair of groups which was calculated through the Spearman’s correlation. * means p < 0.05 FDR corrected.

### The relationship between functional connectivity and levels of consciousness

Furthermore, the mean functional connectivity z-value among the five regions was significantly correlated with levels of consciousness states in the sleep group (ρ = 0.69, p < 0.01), DOC group (ρ = 0.79, p < 0.01), and REM-sleep group (ρ = 0.48, p < 0.01). Similar as the mean strength of functional connectivity, if the functional connectivity with a z-value (> 0.3095) was regarded as an edge, the total edge number among the five brain regions also showed a significant correlation in the sleep group (ρ = 0.78, p < 0.01), DOC group (ρ = 0.81, p < 0.01), and REM-sleep group (ρ = 0.59, p < 0.01) (Fig. 5 and Supplementary fig. 5). All p values above were FDR corrected.

**Fig. 5.**
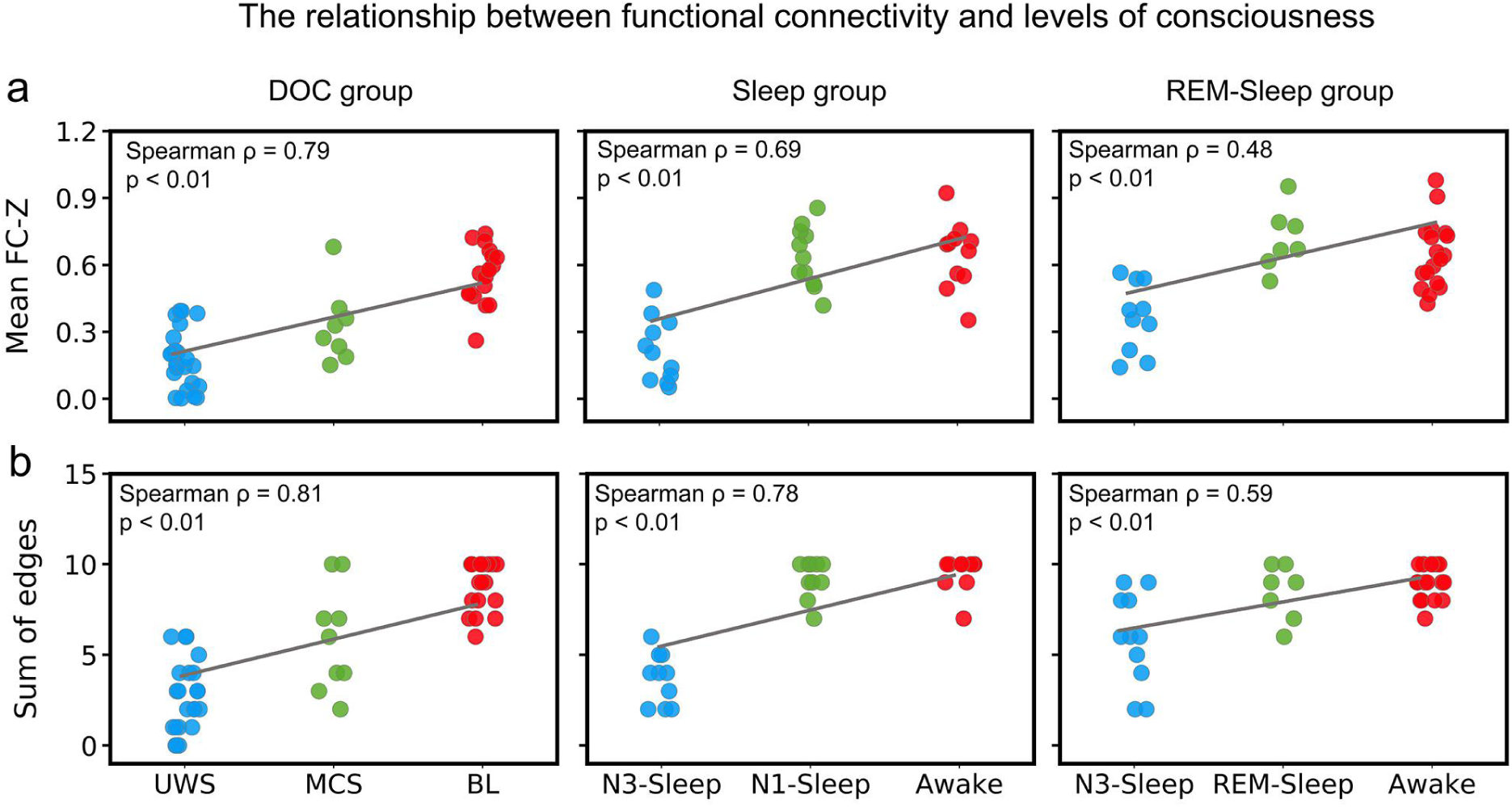
The relationship between functional connectivity and levels of consciousness. **a** The correlation between mean functional connectivity (FC) z-values (of all FCs among the five ROIs: SACC, SMA, LSMG, RSMG and LMTG) for each subject and consciousness states, in patients with DOC (left panel), sleep group (middle panel), and REM-sleep (right panel). **b** The correlation between sum of edges (of all FCs among the five ROIs: SACC, SMA, LSMG, RSMG and LMTG) and consciousness states, in patients with DOC (left panel), sleep group (middle panel), and REM-sleep group (right panel). Edge = one FC with Z value > 0.3095.

### Reduced functional connectivity during unconscious states

Finally, comparing the functional connectivity of each pair regions between consciousness and unconsciousness states, the functional connectivity between SMA and bilateral SMG, the functional connectivity between SACC and LSMG, the functional connectivity between SACC and LMTG, as well as the functional connectivity between bilateral SMG, showed significant reduction in unconsciousness state for all groups (sleep group: awake vs. N3-sleep, N1-sleep vs. N3-sleep; DOC group: BL vs. UWS, MCS vs. UWS; anesthesia group: consciousness vs. anesthesia; especially for REM-sleep group: awake vs. N3-sleep, REM-sleep vs. N3-sleep) (Fig. 6). This result was further confirmed by voxel-wise analysis using the five ROIs (SMA, SACC, bilateral SMG and LMTG) as seed (Supplementary fig. 6).

**Fig. 6.**
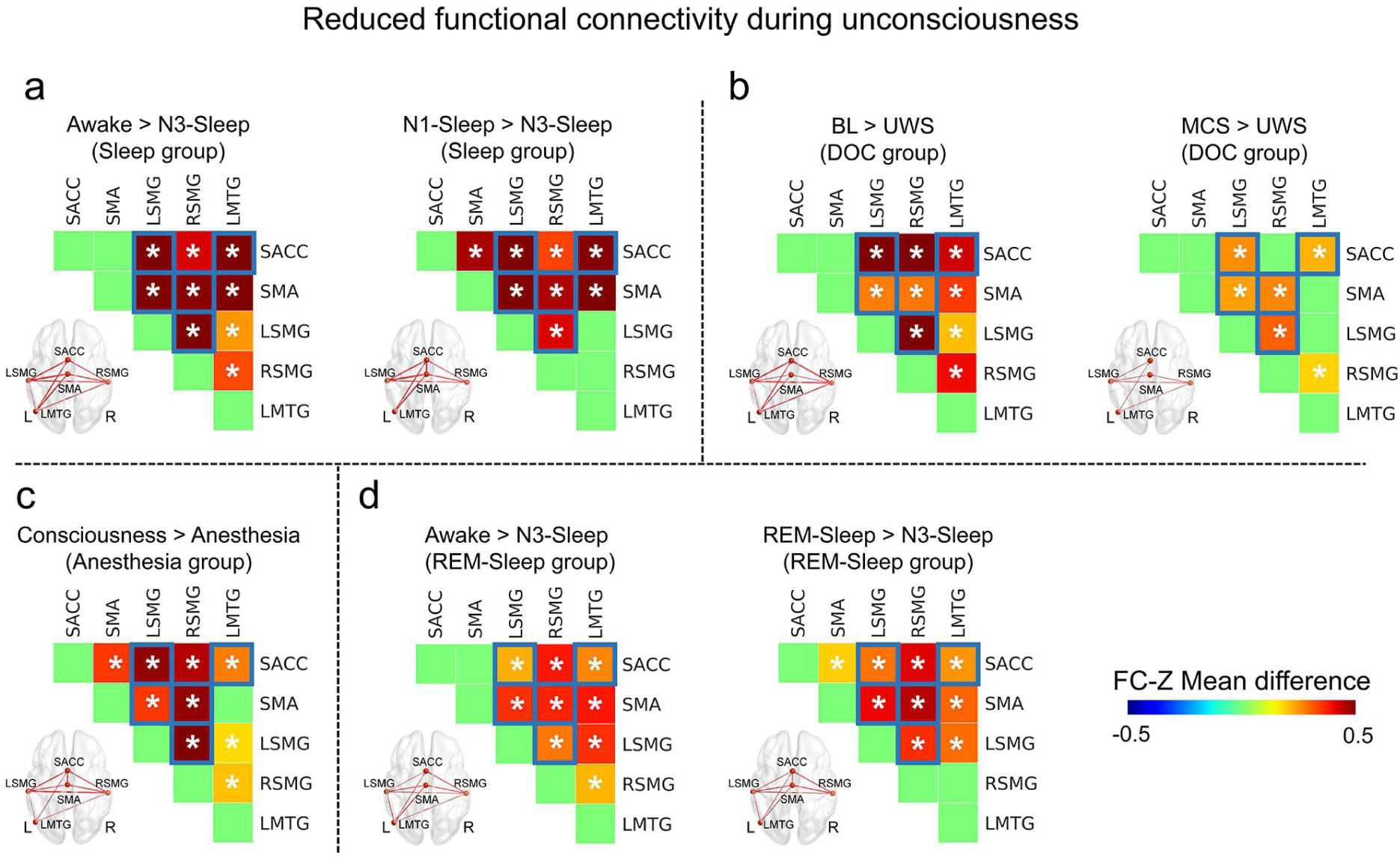
Reduced functional connectivity during unconsciousness. **a** Functional connectivity (FC) with significant reduction in the N3-sleep compared with awake and N1-sleep in the sleep group. **b** FC with significant reduction in UWS compared with BL and MCS in the DOC group. **c** FC with significant reduction in anesthesia compared with consciousness in the anesthesia group. **d** FC with significant reduction in N3-sleep compared with awake and awake in the REM-sleep group. * means p < 0.05, corrected. The functional connectivity marked with blue squares showed consistent reduction during unconsciousness in all four groups. The thickness in brain image represented the difference of functional connectivity between two states.

Additionally, within the above the functional connectivity with significant reduction during unconsciousness, the functional connectivity between SACC and LSMG, the functional connectivity between SACC and LMTG, as well as the functional connectivity between bilateral SMG, were significantly reduced in MCS compared with BL in the DOC group. But there was no functional connectivity which showed significant reduction in the N1-sleep or REM-sleep compared with awake state (Supplementary fig.7).

## Discussion

In this study, we employed a graph-theoretical measure, degree centrality, and functional connectivity analysis to investigate the hubs in the brain’s resting state that support consciousness. Degree centrality reflects the connections of one node with all other nodes throughout the brain. Our results revealed reduced degree centrality in SMA, SACC, bilateral SMG and LMTG in the unconscious states (N3-sleep, anesthesia, and UWS) compared with conscious and altered/partially conscious states (N1-sleep, REM-sleep and MCS). Furthermore, functional connectivity analysis demonstrated that the connections between these five regions were significantly reduced during these unconscious states. Most interestingly, we found that this higher-order sensorimotor integration circuit was fully present in REM-sleep, showing no difference compared with awake states while exhibiting significant difference compared with N3-sleep. This excludes sensory and motor input processing (as both are minimum in REM-sleep) as possible confounds of the higher-order sensorimotor integration circuit as neural basis of consciousness. Therefore, we conclude that regions forming a higher-order sensorimotor integration circuit are central in supporting consciousness through their role as hubs within the brain’s global functional network.

During the REM-sleep, one loses the ability of movement and sensory inputs from the external world. Most interestingly, similar as the REM-sleep, one type of patients named as cognitive motor dissociation (CMD) (Schiff, 2015), shows signs of consciousness with command-following brain activation but no detectable behavioral response (Monti et al., 2010). In the previous literatures, using motor imagery tasks, task-related activation in SMA can be used to identify CMD patients (Owen et al., 2006). However, the difficulty of such tasks and the cooperation of patients could lead to false negatives in the detection of CMD (Pan et al., 2020). Although the brain computer interface (BCI) provides another way to detect the CMD patients by adopting face detection (Pan et al., 2020), the current results may provide a much easier way to identify the CMD patients by using resting-state fMRI dataset without including any active tasks.

Furthermore, previous studies demonstrated global changes of degree centrality during unconsciousness (Hashmi et al., 2017; Tagliazucchi, von Wegner, et al., 2013). Our results extend these findings by showing significantly decreased degree centrality in specific brain regions: the SMA, SACC, bilateral SMG, LMTG in unconscious states compared with conscious and altered conscious states across all groups (N3-sleep compared with N1-sleep, awake, REM-sleep; anesthesia compared with consciousness; and UWS compared with MCS and BL). All these five regions showed high degree centrality in the healthy conscious brain in previous studies (van den Heuvel & Sporns, 2013). Regions with high degree centrality enable efficient communication, and therefore information integration in a global way (van den Heuvel & Sporns, 2013). Consequently, the reduction in the degree centrality in these regions during unconsciousness possibly indicates their role in supporting consciousness within the brain’s global functional network.

Our findings on the relationship between SMA and conscious processing are supported by previous studies in both fully conscious and unconscious subjects. The SMA has been shown to be involved in conscious processing in a variety of different tasks all requiring sensorimotor integration. For instance, one study found that suppressing brain activity in SMA affected visual perception (Martin-Signes, Perez-Serrano, & Chica, 2019). Using the subject’s own name as stimulus, MCS patients showed stronger SMA response to their own names than UWS patients (Qin et al., 2010). SACC is another region involved in motor initiation, in which damages can result in akinetic mutism (Boly et al., 2017). SACC is also a core region of the salience network (Seeley et al., 2007), which shows altered activation during anesthesia (Huang et al., 2014), sleep (Mitra, Snyder, Tagliazucchi, Laufs, & Raichle, 2015), and brain injury (Mitra et al., 2015; Qin et al., 2015). Our previous study showed the functional connectivity between SACC and insula was correlated with the consciousness levels in DOC patients (Qin et al., 2015). This was consistent with the voxel-wise functional connectivity in the current study (Supplementary fig. 6).

Beside these motor regions, the SMG, a higher-order sensory region, showed significant reduction of global integration during unconsciousness (Luppi et al., 2019). More importantly, the SMG, is part of the inferior parietal lobule (IPL) which was reported to show reduced brain activation in unconscious states (Lichtner et al., 2018). Since the IPL closes to the temporal and occipital cortices and receives both auditory and visual information (Dehaene & Changeux, 2011) as well as heartbeat awareness and body-centered perception (e.g., face and body) (Blanke, Slater, & Serino, 2015), it is suggested to play a central role in integrating external and internal sensory information. Similar as SMG, LMTG was regarded as a part of temporo-parietal-occipital hot zone, and was more related to content-specific neural correlates of consciousness in previous studies (Koch et al., 2016). Furthermore, LMTG was also involved in the visual word detection task (Dehaene et al., 2001), and in dreaming experience compared with no dreaming (Siclari et al., 2017).

More interestingly, the current results showed that the sum of functional connectivity (mean strength and edge) among these five regions was correlated with the levels of consciousness. This strongly suggests that they could form a circuit within the brain’s global functional networks whose higher-order sensorimotor integration is key in supporting consciousness. Taken together, the reduced connectivity of these five regions suggests that loss of consciousness may be contributed by a reduction in multiple high-order functional integrations, especially the integrated sensory and motor information, i.e., sensorimotor integration, which consistently showed reduction in all the unconscious state in the four groups.

The Default-mode network also showed reduced degree centrality several contrasts between conscious/altered conscious and unconscious states, such as UWS (compared with MCS), as well as anesthesia (compared with consciousness in anesthesia group), but not in all the contrasts between conscious/altered and unconscious states (Fig.2). This was consistent with the previous studies. Some of them showed that the functional connectivity was correlated with levels of consciousness (Vanhaudenhuyse et al., 2010), while some of them indicated that default-mode network was related to the recovery of consciousness, not the levels of consciousness (Norton et al., 2012).

Several issues should be noted. First, deep, surgical-level anesthesia was used, thus some of the anesthetic effects on the BOLD signal could be physiological in nature and unrelated to unconsciousness. In the current study, during propofol-induced anesthesia in 17 participants, there was a significant reduction in the degree centrality in SMA, SACC, bilateral SMG and LMTG during unconsciousness. Although there were only six participants receiving sevoflurane, their results also showed similar trends in all these regions, and significant degree centrality reduction in right SMG (See Supplementary fig. 8). Hence, our results suggest that there are no major differences in degree centrality and functional connectivity between the two drugs. In the future, it may be useful to test other anesthetic drugs with substantially different pharmacology (e.g., ketamine) to confirm the current results. The second issue is that we did not specifically investigate dreams during both NREM and REM but took REM-sleep simply as an index of the presence of consciousness (Windt, Nielsen, & Thompson, 2016). Although different from wakefulness, the REM-sleep is considered a particular form of environmentally “disconnected” consciousness (Tagliazucchi, Behrens, & Laufs, 2013). Here we not only showed the similarity of functional connectivity of the five brain hubs between a presumed dream state, i.e., REM, and the fully awake conscious state, but also its difference from that in N3-sleep. Finally, although the results about the REM-sleep group were consistent with other groups, and supported by the previous literatures, the sample size with seven participants was relatively small. In the future, more studies should be carried out to further confirm the current results.

In summary, we demonstrated that SMA, SACC, bilateral SMG and LMTG participate in forming a higher-order sensorimotor integration circuit within the brain’s global functional network that supports consciousness. All these regions showed significant decreases in degree centrality and functional connectivity with each other during different unconscious conditions, i.e., anesthesia, N3-sleep, and patients with UWS. Furthermore, the mean strength and edge of functional connectivity of the five regions were significantly correlated with levels of consciousness. Interestingly, these properties were unchanged from wakefulness during the REM-sleep. Based on their known functions in movement initiation and sensory information processing, we conclude that the here identified higher-order sensorimotor integration circuit is essential for maintaining wake-like consciousness.

## Materials and Methods

### Participants and data acquisition – Sleep group

We recruited 12 healthy men with regular sleep duration of 7–8 h per night and consistent bed/wake time for at least 4 days. They had no daytime nap habits, no excessive daytime sleepiness, and no history of neurological or psychiatric disorders. Participants’ age ranged from 20 to 27 years (mean ± SD = 23 ± 2.5 years). The participants were requested not to consume alcohol or caffeine-containing food or drinks on the day of the experiment. Before scanning, we requested the participants to complete the Pittsburgh Sleep Quality Index (PSQI), including sleep quality, duration, and efficiency, to characterize the general quality of their sleep within the previous month. Informed consent was obtained from all participants prior to the experiments in accordance with the protocol approved by the Institutional Review Board of National Yang-Ming University.

We conducted simultaneous EEG–fMRI recordings for the sleep protocol. EEG was recorded using a 32-channel MR-compatible system (Brain Products, Gilching, Germany). The 32 electrodes, including 30 EEG channels, one electrooculography (EOG) channel, and one electrocardiogram (ECG) channel, were positioned according to the international 10/20 system. The electrode-skin impedance was reduced to <5 kΩ using abrasive electrode paste (ABRALYT HiCl). The EEG signal was synchronized with the MR trigger and recorded using Brain Vision Recorder software (Brain Products) with a 5 kHz sampling rate and a 0.1 μV voltage resolution. Low-pass and high-pass filters were set at 250 Hz and 0.0159 Hz, respectively, with an additional 60-Hz notch filter.

MRI data were acquired using a 3T Siemens Tim Trio system (Erlangen, Germany). High-resolution T1-weighted anatomical images were acquired prior to functional scans for geometric localization. Head motion was minimized using customized cushions. Functional scans during sleep were subsequently acquired using a T2*-weighted EPI sequence (TR/TE/θ = 2500 ms/ 30 ms/80°, field of view = 220 mm, matrix size = 64 × 64, 35 slices with 3.4 mm thickness, maximum of 3000 scans) aligned along the AC–PC line, allowing whole-brain coverage. The sleep protocol was conducted at midnight and the participants were instructed to try to fall asleep after the EPI scan started. Other details of experimental and preprocessing procedures can be found in previously published work (Tsai et al., 2014). A licensed sleep technician in Kaohsiung Medical University Hospital visually scored the sleep stages for every 30 s epoch, according to the current criteria of the American Academy of Sleep Medicine (AASM). In the current study, awake state (before sleep), N1-sleep, and N3-sleep were included in the analysis. Since the Sleep group did not include the REM-sleep stage datasets we included another REM-Sleep group which included a REM-sleep dataset to control the effects of immobility and lack of sensory inputs from the environment for most unconscious states, which in the current study were the N3-sleep, anesthesia and MCS.

### Participants and data acquisition – REM-Sleep group

These datasets were from a previously published study (Fang, Ray, Owen, & Fogel, 2019) using simultaneous EEG–fMRI recordings. There were 24 healthy participants (age from 18 years to 34 years, mean ± SD = 23.8 ± 4.1 years, 9 female) included in the current analysis. The awake-state was recorded in 19 participants (which occurred during sleep, not before sleep), the N3-state was recorded in 11 participants and the REM-sleep was recorded in seven participants. One participants had all of the awake, N3-sleep and REM-sleep states; Two participants had both N3-sleep and REM-sleep; Two participant had both awake and REM-sleep. Seven participants had awake and N3-sleep. Each sleep state had a minimum of 90 volumes. The sleep stages were scored in the same way as sleep group 1. See the previous study for detailed information about the EEG and fMRI data acquisition parameters and data preprocessing (Fang et al., 2019).

### Participants and data acquisition – Anesthesia group

Seventeen participants received intravenous propofol anesthesia and six subjects received inhalational sevoflurane anesthesia. The participants’ age ranged from 26 years to 63 years (mean ± SD = 45.8 ± 11.8 years, 11 female). All the subjects attended elective transsphenoidal approach for resection of pituitary microadenoma. Informed written consent was obtained from each subject. The study was approved by the Ethics Committee of Shanghai Huashan Hospital, Fudan University, Shanghai, China.

For the propofol group, we achieved a 3.0-5.0 μg/ml plasma concentration by using a target-controlled infusion (TCI) based on the Marsh model. This was followed by remifentanil (1.0 μg/kg) and succinylcholine (1 mg/kg) to facilitate endotracheal intubation. We then maintained the TCI propofol at a stable effect-site concentration (4.0 μg/ml) which reliably induced an unconscious state in the subjects. For the sevoflurane group, we gave the subjects 8% sevoflurane in 100% oxygen, adjusting fresh gas flow to 6 L/min, combined with remifentanil (1.0 μg/kg) and succinylcholine (1.0 mg/kg). This was maintained with 2.6% (1.3 MAC) ETsevo in 100% oxygen, and a fresh gas flow at 2.0 L/min. The concentration of sevoflurane successfully maintained a loss of consciousness in subjects, classified as ASA physical status I or II. The anesthetic effects on the brain are considered to be solely pertaining to propofol and sevoflurane because of the quick elimination of the analgesic remifentanil and depolarized neuromuscular relaxant succinylcholine from plasma. To confirm the effects of each drug, the results for each drug were also presented separately in the supplementary materials.

During the anesthetic state, subjects were given intermittent positive pressure ventilation, with tidal volume at 8-10 ml/kg, respiratory rate at 10-12 beats per minute, and PetCO_2_ (end tidal partial pressure of CO_2_) at 35-37 mmHg. All subjects fulfilled the criteria of deep sedation: neither a response to verbal commands (“squeeze my hand”), nor response to prodding or shaking was observed during anesthesia, corresponding to Ramsay 5–6 and an OAA/S score of 1. In addition, no subject reported explicit memory in the post-operative follow-up. Therefore, all subjects were considered unconscious during anesthesia.

All the datasets had the same fMRI acquisition parameters. A Siemens 3T scanner used T2*-weighted EPI sequence to acquire functional images of the whole brain (TR/TE/θ= 2000ms/30 ms/90°, FOV = 192 mm, matrix size = 64 × 64, 25 slices with 6-mm thickness, gap = 0 mm, 240 scans). A high-resolution T1-weighted anatomical image was also acquired for all participants. EPI data acquisition was conducted both in wakefulness prior to anesthesia and in the anaesthetized state. The subjects were instructed to relax and keep their eyes closed during scanning. Following this, subjects were anaesthetized and given full hydration with hydroxyethyl starch to avoid hypotension. Fifteen minutes after stabilization of anaesthetic levels and hemodynamics, the anaesthetized state resting-state fMRI scan was done.

### Participants and data acquisition –patients with disorders of consciousness

To minimize the effects of structural distortion and maintain local neural integrity on functional connectivity, we included 50 structurally preserved brain injury patients in the current study. These structurally well-preserved patients, which were visually selected by author XW and then checked by author PQ, had limited brain lesions and limited brain structure distortion (Supplementary Table 1 for detailed demographic and clinical information, and Supplementary fig. 1 for each patient’s structural image). The UWS and MCS patients were assessed using the Coma Recovery Scale-Revised (CRS-R) (Giacino, Kalmar, & Whyte, 2004) before fMRI scanning (T0). The Glasgow Outcome Scale (GOS) was carried out at least three months after the scan session (T1). Additionally, we included 52 healthy participants (ages ranged from 23 to 59 years, 15 females) from one of our previous studies (Qin et al., 2015). Informed written consent was obtained from the patients’ legal representatives. The study was approved by the Ethics Committee of Shanghai Huashan Hospital, Fudan University, Shanghai, China.

For the datasets in the current study, the MR images were acquired on the same Siemens 3 Tesla scanner. Functional images were acquired using a T2*-weighted EPI sequence (TR/TE/θ = 2000 ms/35 ms/90°, FOV = 256 x 256 mm, matrix= 64 x 64, 33 slices with 4-mm thickness, gap = 0 mm, 200 scans). Each volume had 33 axial slices, covering the whole brain. Two hundred volumes were acquired during rest. A high-resolution T1-weighted anatomical image was acquired for all participants for functional image registration and localization. The patients were instructed to take a comfortable supine position, relax, close their eyes, and not concentrate on anything in particular during the scanning. All participants wore earplugs to minimize the noise of the scanner.

### Data preprocessing

Anatomical images from the two sleep groups, anesthesia, and patient with DOC were segmented into grey matter (GM), white matter (WM), and cerebrospinal fluid (CSF), using the FAST tool from the FSL software package (http://www.fmrib.ox.ac.uk/fsl/). Functional images were processed using the AFNI software package. After discarding the first two volumes, functional images underwent a preprocessing procedure which included: slice-timing correction; two- and three-dimensional head-motion corrections; masking for the removal of the skull; and spatial smoothing using an 8-mm kernel. Time-series were then intensity normalized by computing the ratio of the signal in each voxel at each time point to the mean across all time points, and then multiplied by 100. The six estimated head motion parameters and the mean time-series from the white matter (WM) and the cerebrospinal fluid (CSF), which were defined using partial volume thresholds of 0.99 for each tissue type, were considered as noise covariates and were regressed out from the data. The data were band-pass filtered preserving signals between 0.01 and 0.08 Hz. Structural and functional images from all the three groups were transformed into MNI standard space (3 x 3 x 3 mm^3^ resolution).

### Head motion correction

For all the datasets, the issue of head motion was rigorously addressed as minor differences in motion have been shown to cause artificial group differences (Power, Barnes, Snyder, Schlaggar, & Petersen, 2012). Motion was quantified as the Euclidean distance calculated from the six rigid-body motion parameters for two consecutive time points (AFNI, 1d_tool.py). Any instance of movement greater than 0.5 mm was considered as excessive motion, for which the respective volume as well as the immediately preceding and subsequent volumes were removed. In order to obtain reliable results, participants with less than 90 scans remaining were excluded (Yan, Craddock, He, & Milham, 2013).

### Degree centrality changes in different conscious levels

For the two sleep groups, anesthesia, structurally well-preserved patients, and healthy controls, voxel-wise degree centrality analysis was performed within a grey matter mask (Yeo et al., 2011) using the AFNI program 3dTcorrMap. In the current analysis, each voxel constitutes a node and each effective functional connection (Pearson correlation at r > 0.3) between the node and any other voxel (node) within the grey matter mask was an edge. Degree centrality was the number of edges connecting to a node. We hereby adopted different thresholds (Pearson correlation at r > 0.2 and r > 0.4) for validation.

For the sleep group, awake and N1-sleep were regarded as consciousness or reduced consciousness states while N3-sleep as an unconscious state (Laureys, 2005); for the anesthesia group, awake consciousness was the conscious state while anesthetic state was an unconscious state. For the DOC group, the BL and MCS were regarded as consciousness or reduced consciousness states while UWS as an unconscious state. In order to identify the common brain regions with degree centrality changes during unconscious states, paired t-tests were performed to compare voxel-wise differences between unconscious and conscious/reduced conscious states for the anesthesia group. For the sleep group, repeated measures ANOVA was applied to analyze the difference of degree centrality values between unconscious and conscious/reduced conscious states with AFNI program (3dMVM). For the DOC group, ANCOVA was applied to analyze the voxel-wise contrasts of degree centrality values through 3dMVM. Age, gender, head motion, the length of time since brain injury, and the length of BOLD signal series were regarded as covariates. All tests above were two-sided. For all these contrasts, a cluster-level significance threshold of p < 0.05 following FWE correction (p < 0.005 uncorrected, cluster size > 60 voxels, 3dClusterSim, AFNI) was used. The common brain regions with degree centrality reduction during unconsciousness detected in the anesthesia group, sleep group and DOC group were defined as the overlapping brain regions (volume > 20 voxels).

For each common brain region with degree centrality reduction during unconsciousness in the sleep group, anesthesia group and DOC group, the coordinates of cluster centers were used to make ROIs (sphere with r = 8mm). Then for the REM-sleep group, mean degree centrality values within these ROIs were computed, respectively. For the REM-sleep group, for each ROI, the Kruskal-Wallis H test was applied to analyze the contrasts of degree centrality values; post-hoc test included: N3-sleep vs. REM-sleep, N3-sleep vs. Awake, and REM-sleep vs. awake. Awake-state and REM-sleep were regarded as consciousness or reduced consciousness states while N3-sleep was regarded as an unconscious state. All tests above were two-sided. The effect size was estimated using *η*^2^. The FDR correction was used to control for the multiple comparison problem across all post-hoc tests among the five brain regions in the REM-sleep group.

### Functional connectivity changes between brain regions with decreased degree centrality during unconscious states

Following the degree centrality analysis, ROI-based functional connectivity was performed for the ROIs from the above analysis which showed degree centrality reduction in unconsciousness. Mean time-series from each region was calculated, and then the Pearson correlation coefficient was computed for each pair of regions. Fisher’s Z transformation was used to transform the correlation r-values to normally distributed Z-values, which were then compared between unconscious and conscious states. In order to determine whether the functional connectivity of all the five regions was significant during consciousness or awake-states within each group, a one-sample t-test was applied to compare Z-values to 0.3095 (r value = 0.3). To test whether the relationship among all the pairs of functional connectivity was consistent during consciousness states, the correlations between the functional connectivity z-values during consciousness or awake states from one group and the values during the consciousness states from other groups were calculated by means of the Spearman’s correlation. The p-value of the correlation coefficient were FDR corrected. Additionally, the relationship between the mean functional connectivity z-value among all the pairs of ROI connections and levels of consciousness states were investigated in the sleep group, DOC group and REM-sleep group using the Spearman’s correlation. Similar as the mean strength of functional connectivity, if the functional connectivity with a z-value (> 0.3095) was regarded as an edge, the relationship between total edge number among the five brain regions and levels of consciousness was also investigated using Spearman’s correlation. The p-value were FDR corrected. Since the anesthesia group only included two levels of consciousness it was not included in this correlation analysis.

Finally, to determine the functional connectivity changes from consciousness to unconsciousness, the Wilcoxon rank-sum test was performed for each pair of ROIs in the anesthesia group. FDR corrections was adopted for all pairs of ROIs in the anesthesia group. For the sleep group, the Friedman’s test was performed to compare the functional connectivity between each pair of ROIs among awake, N1-sleep, and N3-sleep. The FDR correction was adopted for all the post-hoc tests in the sleep group. For patients with DOC, the Kruskal-Wallis H test was applied to analyze the functional connectivity between each pair of ROIs among the UWS, MCS, and BL. The FDR correction was adopted for all the post-hoc tests in the DOC group. For the REM-sleep group, the Kruskal-Wallis H test was also applied to analyze functional connectivity between each pair of ROIs in the N3-sleep, REM-sleep and awake. Furthermore, the post-hoc test p-values of the REM-sleep group were corrected using the FDR correction. All tests above were two-sided. For all the statistics, degree centrality values or functional connectivity values beyond the range (mean±2SD) of each group were excluded.

## Acknowledgments

We are also grateful to Dr Junrong Han and Ms. Mingxia Wang for their help in writing the manuscript.

## Funding

This work was sponsored by grants from Key Realm R&D Program of Guangzhou (202007030005 to PQ), the National Natural Science Foundation of China (Grant 31771249 to PQ; Grants 81571025 to XW), International Cooperation Project from Shanghai Science Foundation (No.18410711300 to Xuehai Wu), Shanghai Science and Technology Development funds (No. 16JC1420100 to Y.M), Shanghai Municipal Science and Technology Major Project (No.2018SHZDZX01 to Y.M), Natural Science Foundation and Major Basic Research Program of Shanghai (16JC1420100), Ministry of Science and Technology in Taiwan (MOST 105-2628-B-038-013-MY3), National Science Foundation of China (31471072), Taiwan Ministry of Science and Technology (105-2410-H-038-006-MY3, 105-2410-H-038-005-MY2); and Taipei Medical University (104-6402-006-110), and National Institute of General Medical Sciences of the National Institutes of Health (R01-GM103894, to AGH), Canadian Institutes of Health Research (CIHR), Michael Smith Foundation (EJLB-CIHR), The Hope for Depression Research Foundation (HDRF). This research project was also supported by the HBP Joint Platform to GN, funded from the European Union’s Horizon 2020 Framework Program for research and Innovation under the specific Grant Agreement No 785907 (Human Brian Project SGA 2).

## Author contributions

All authors contributed to the critical revision and final approval of the manuscript version for publication. PQ, XW, YM and GN designed this study and wrote the manuscript. PQ and HW performed the data analysis. CW, XW, JZ, WT, ZH, SF and YZ also contributed to the data acquisition. ND, XW, AGH, SF and TL also contributed to writing of the manuscript.

## Competing interests

The authors report no competing interests.

## Data and materials availability

All data needed to evaluate the conclusions in the paper are present in the paper and the Supplementary Materials. Additional data related to this paper may be requested from the corresponding author.

## Supplementary information

**Supplementary Table 1.**
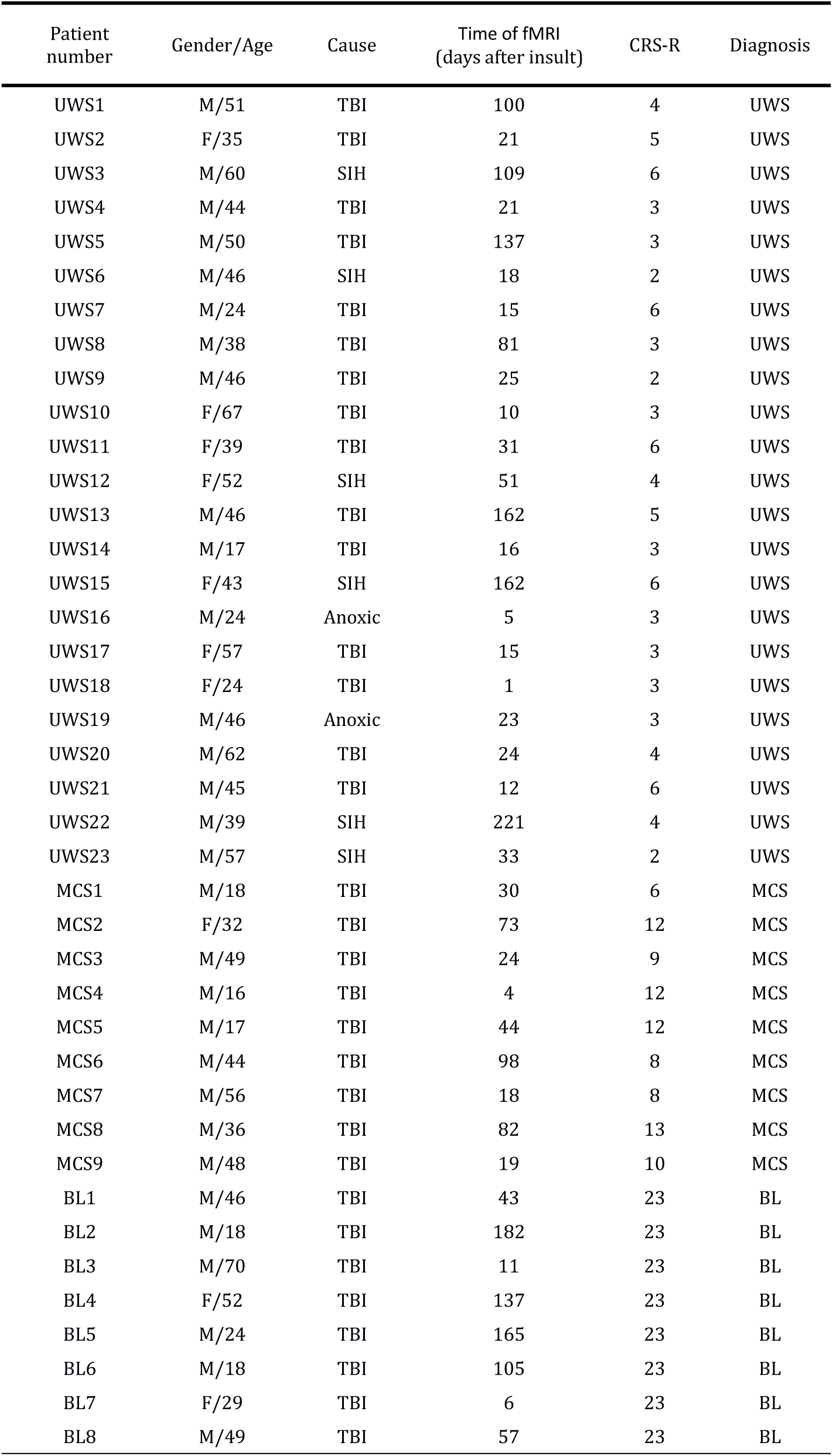
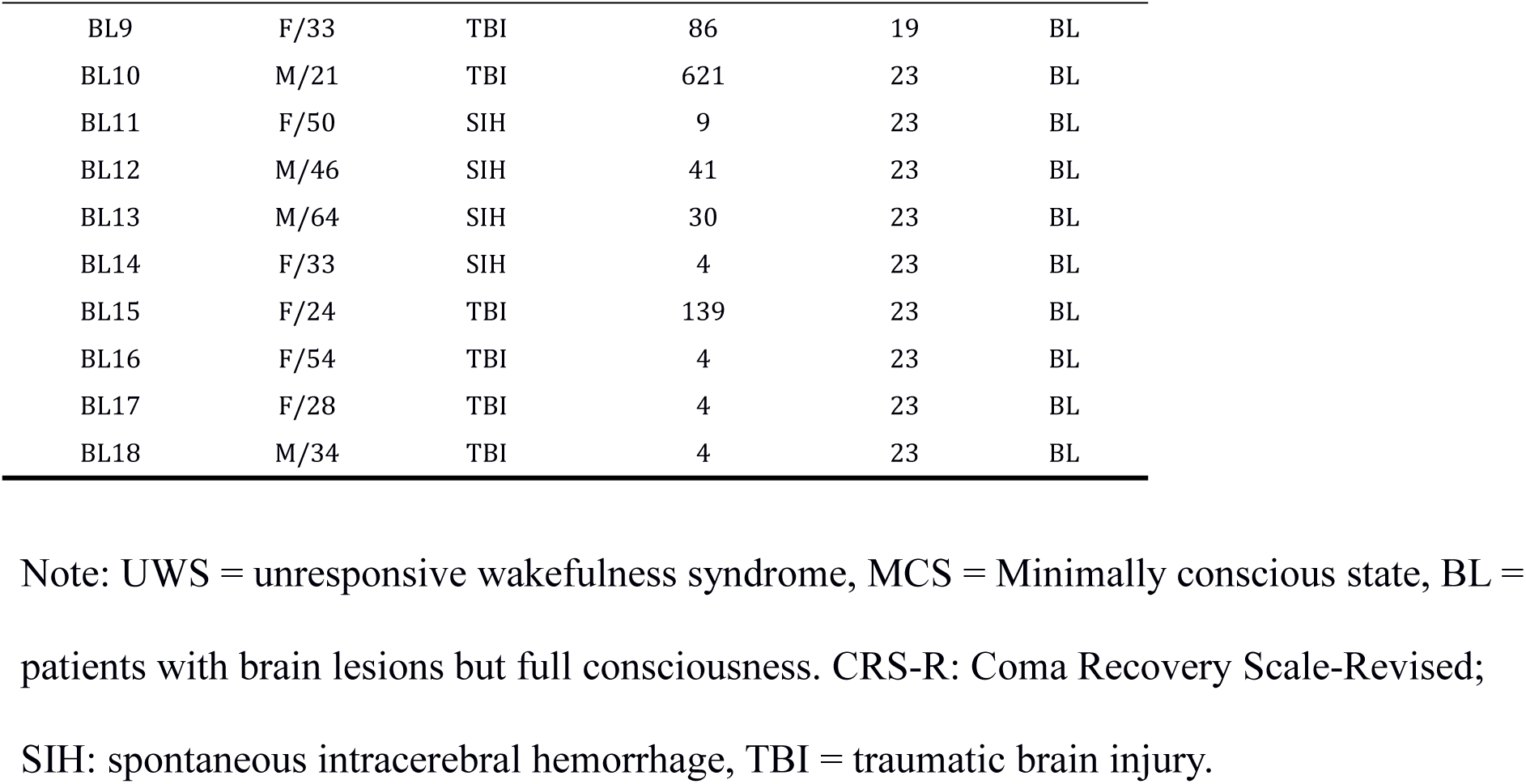
Demographic and clinical information for patients with disorders of consciousness

**Supplementary Table 2.** Coordinates for global hubs with reduced degree centrality in unconsciousness

| Brain regions | Lateral | Coordinates (MNI) |  |  |
| --- | --- | --- | --- | --- |
|  |  | X | Y | Z |
| SACC | M | -2 | 6 | 41 |
| SMA | M | -2 | -14 | 52 |
| SMG | L | -59 | -24 | 21 |
|  | R | 56 | -26 | 24 |
| MTG | L | -54 | -68 | 7 |

Note: The overlapped regions showing significantly reduced voxel-wise DC during all kinds of unconscious states (N3-sleep, anesthesia and unresponsive wakefulness syndrome). SACC = supragenual anterior cingulate cortex; SMA = supplementary motor area; SMG = supramarginal gyrus; LMTG = left middle temporal gyrus.

**Supplementary fig. 1:**
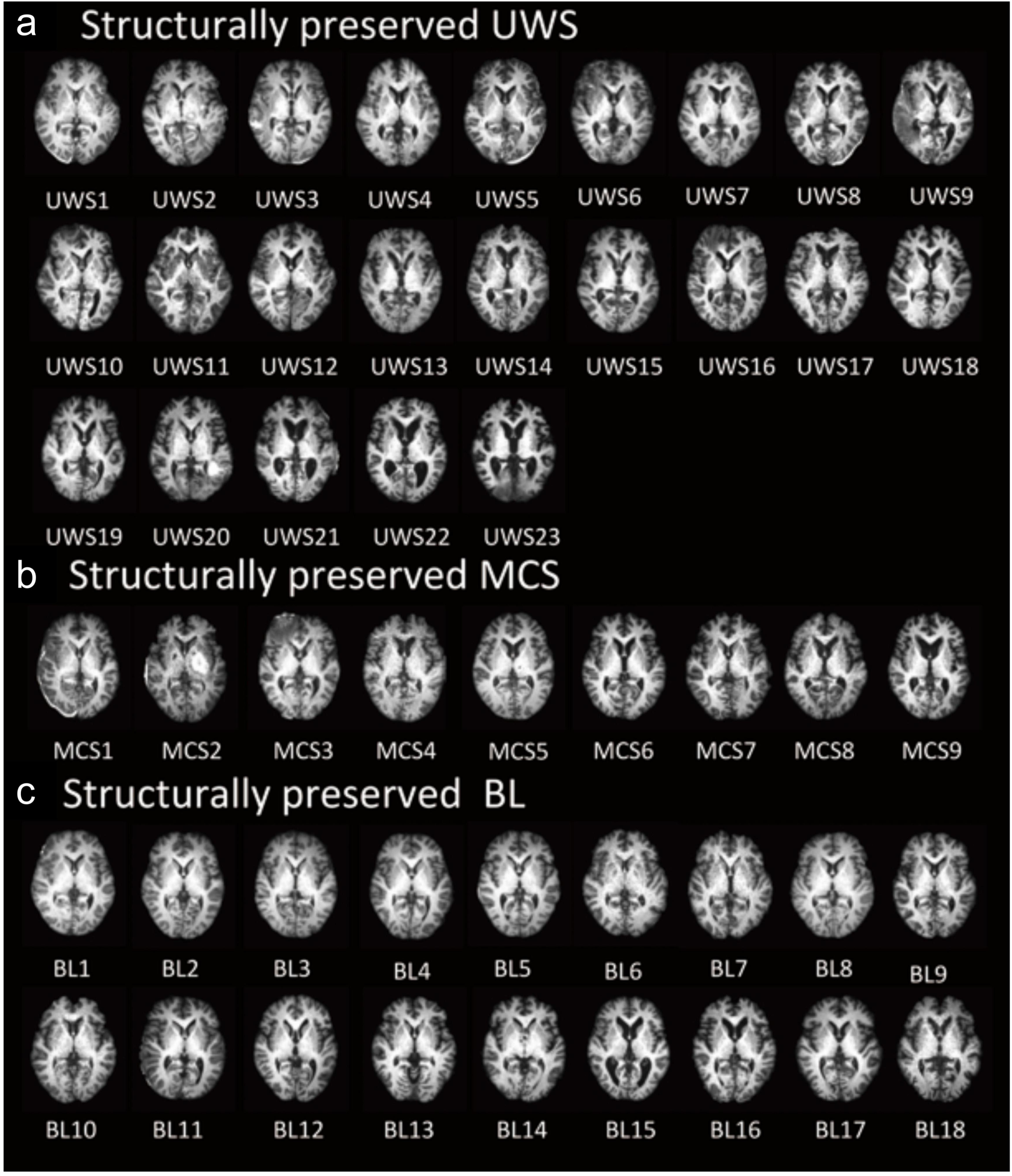
Structure Image patients with disorders of consciousness. (a) UWS = unresponsive wakefulness syndrome, (b) MCS = minimally conscious state, (c) BL = patients with brain lesions but full consciousness.

**Supplementary fig. 2.**
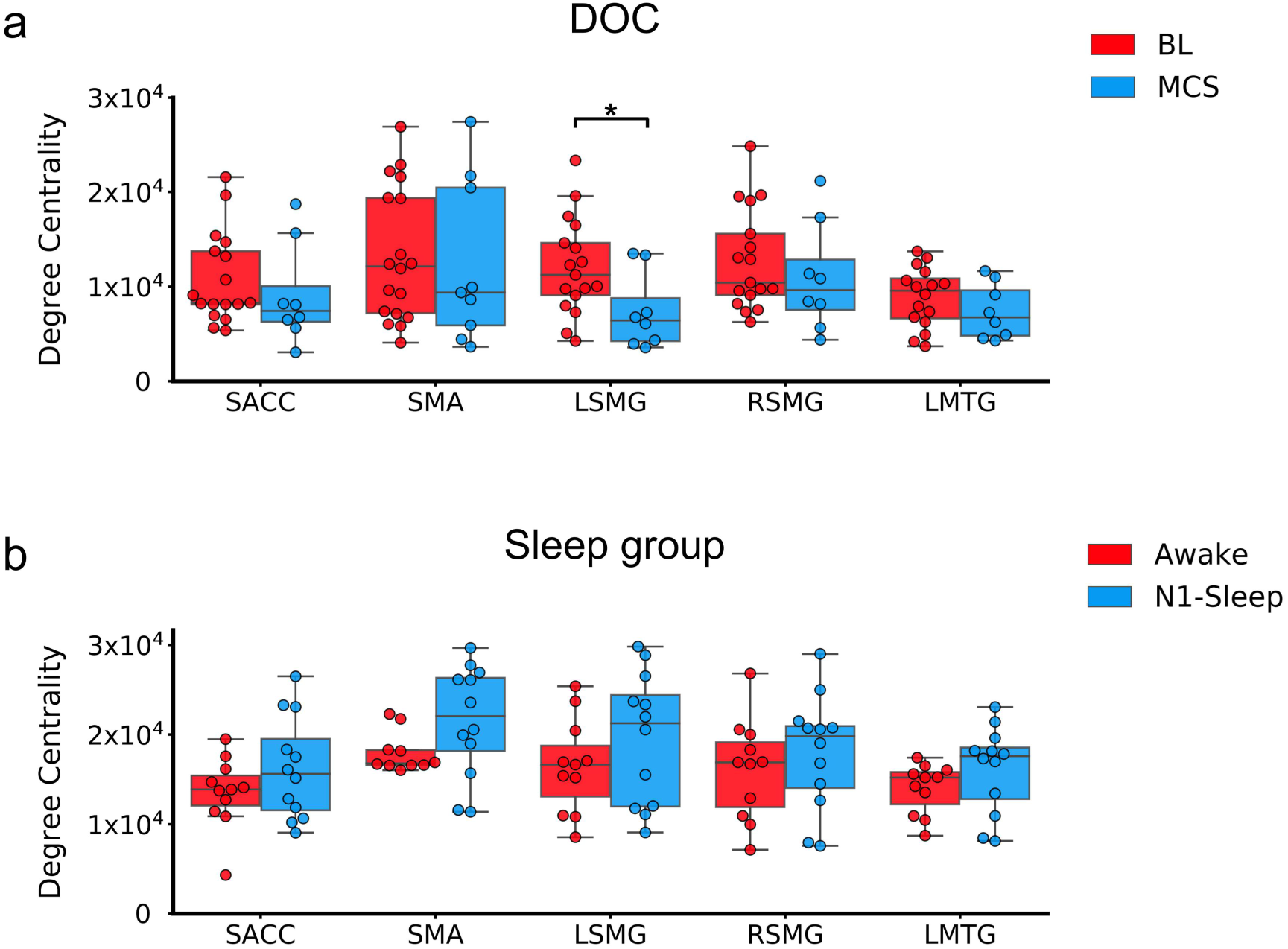
The degree centrality in DOC and sleep group. (a) The degree centrality in BL and MCS in DOC group. (b) The degree centrality in Awake and N1-sleep. SACC = supragenual anterior cingulate cortex; SMA = supplementary motor area; LSMG = left supramarginal gyrus; RSMG = right supramarginal gyrus; LMTG = left middle temporal gyrus. * means p < 0.05 FDR corrected.

**Supplementary fig. 3.**
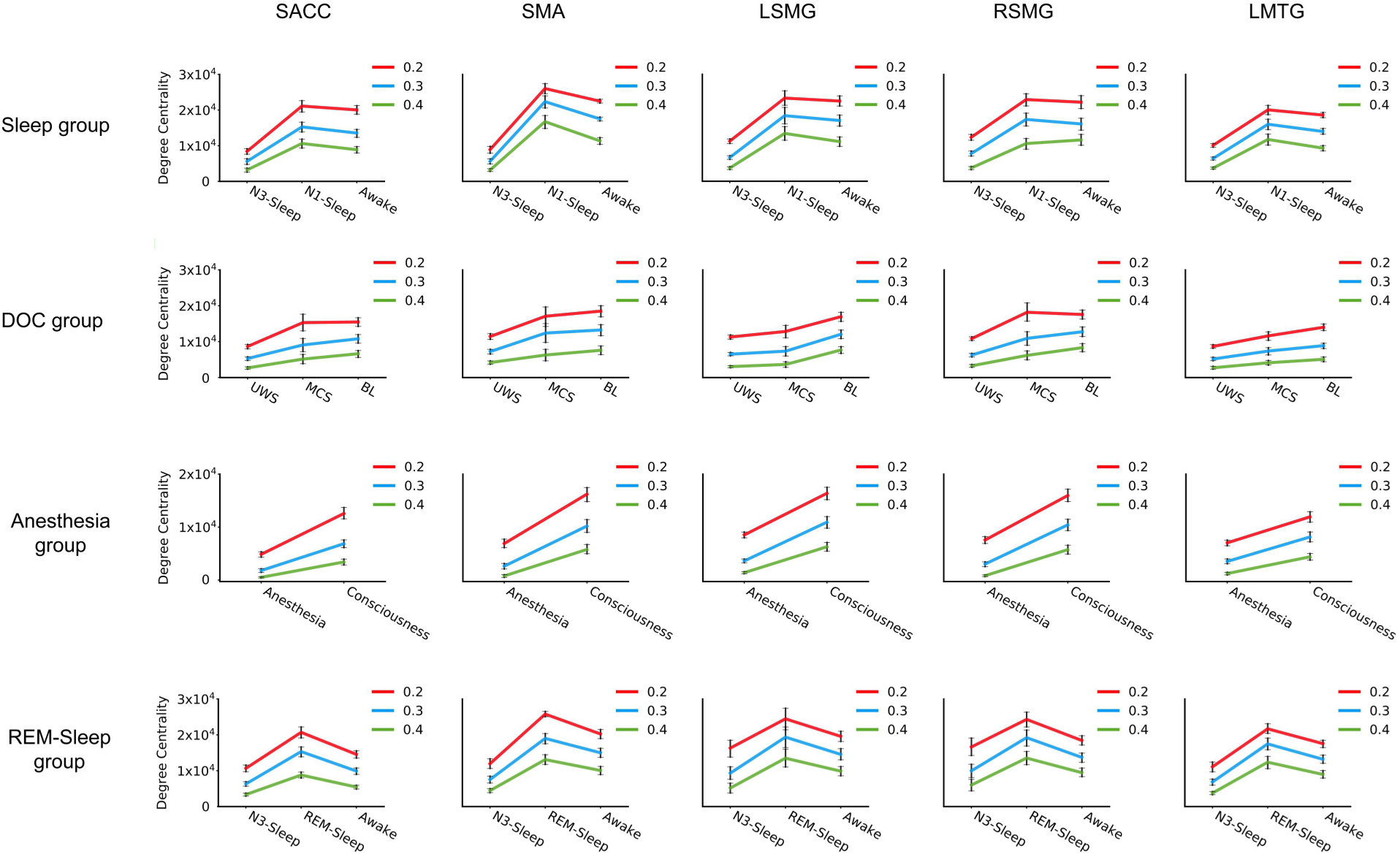
The degree centrality values (with different thresholds) in global hubs. 0.3 means r > 0.3; 0.2 means r > 0.2; 0.4 means r > 0.4. SACC = supragenual anterior cingulate cortex; SMA = supplementary motor area; LSMG = left supramarginal gyrus; RSMG = right supramarginal gyrus; LMTG = left middle temporal gyrus.

**Supplementary fig. 4.**
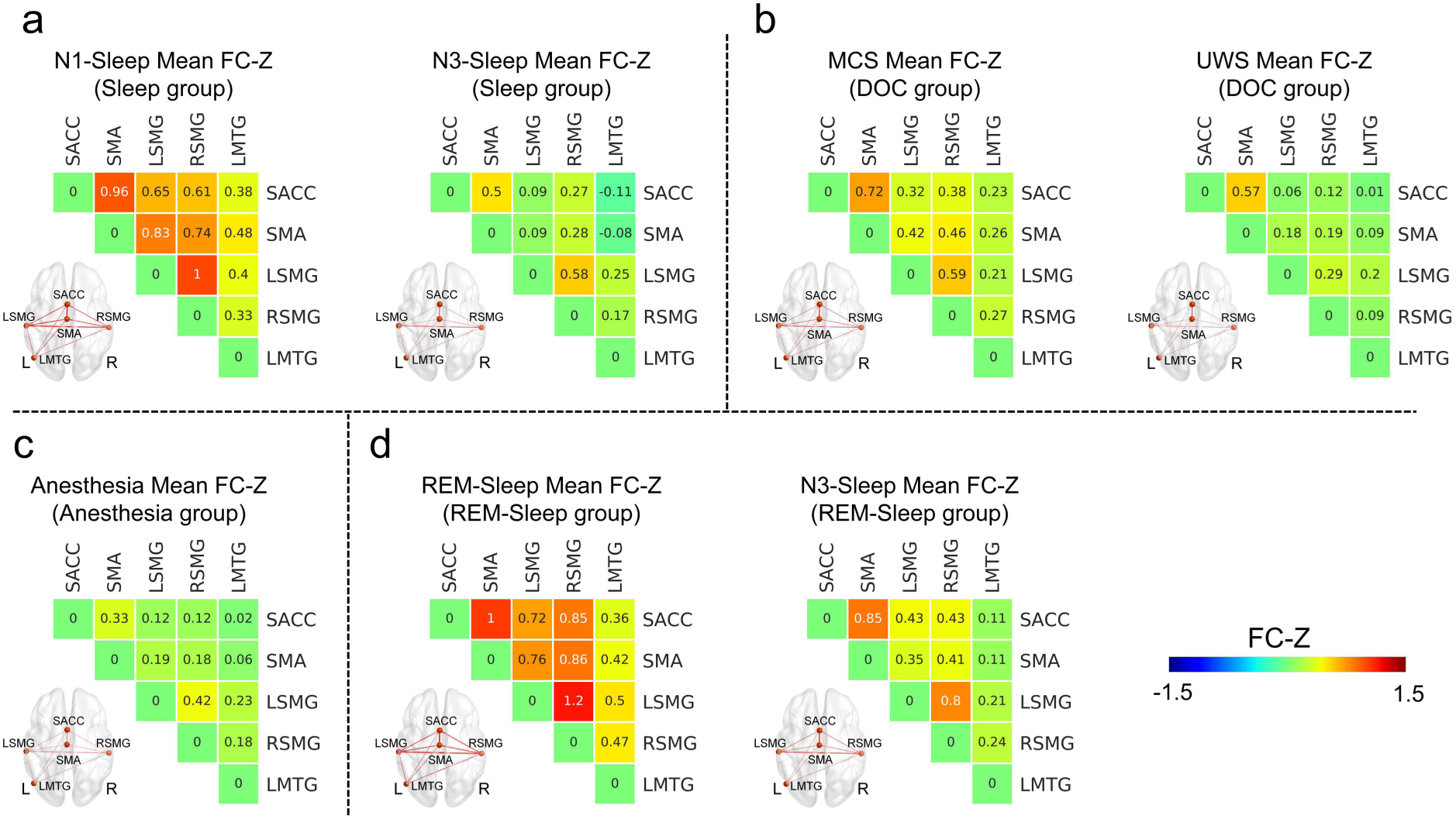
ROI-based functional connectivity in altered conscious states. (a) The mean of functional connectivity z-values of N1-sleep and N3-sleep states in sleep group. (b) The mean of functional connectivity z-values of MCS and UWS in DOC group. (c) The mean of functional connectivity z-values of anesthesia state. (d) The mean of functional connectivity z-values of REM-sleep and N3-sleep states in REM-sleep group. SACC = supragenual anterior cingulate cortex; SMA = supplementary motor area; LSMG = left supramarginal gyrus; RSMG = right supramarginal gyrus; LMTG = left middle temporal gyrus.

**Supplementary fig. 5.**
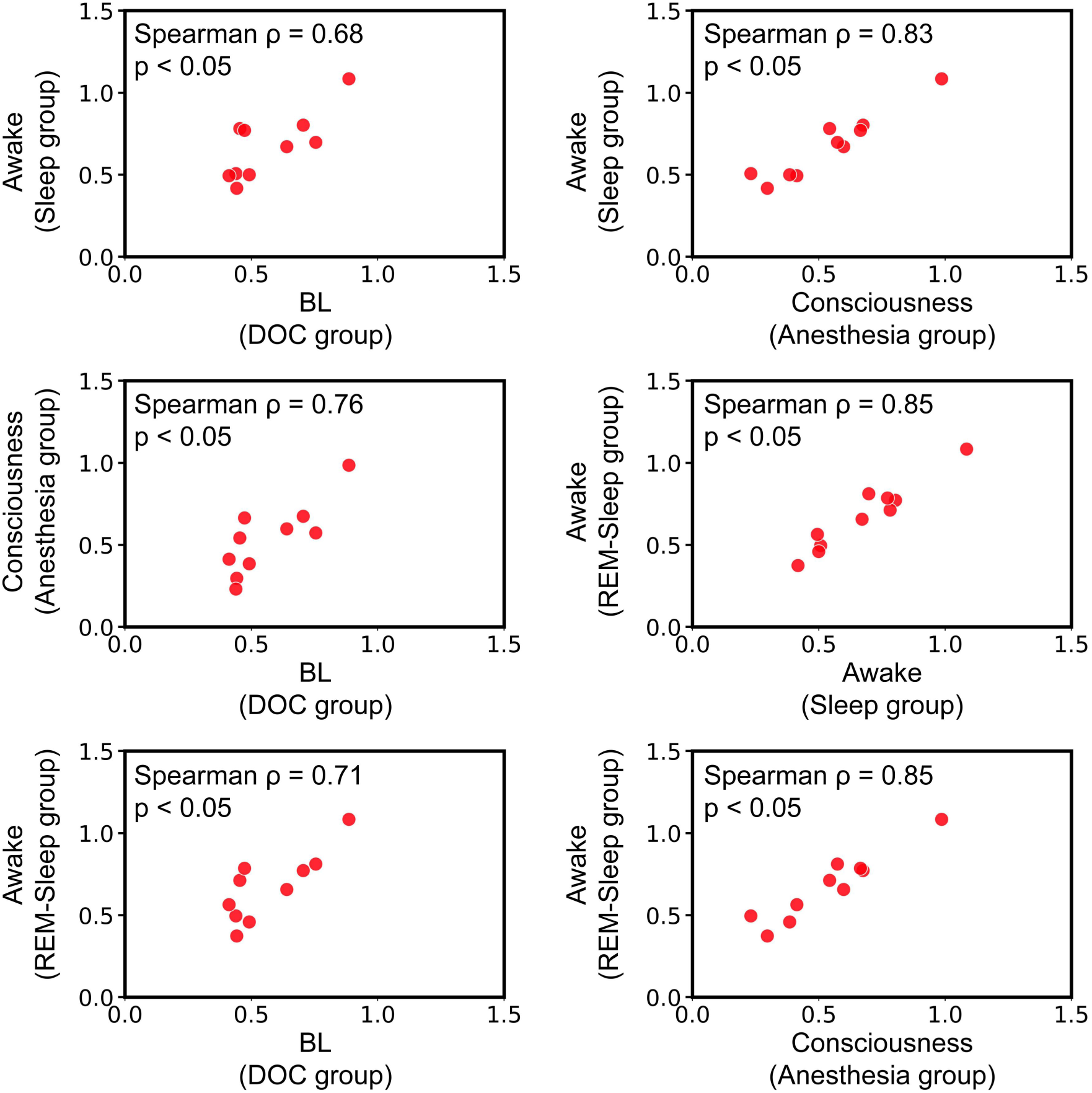
The relationship of functional connectivity (FC) between each of pair of groups during conscious states.

**Supplementary fig. 6.**
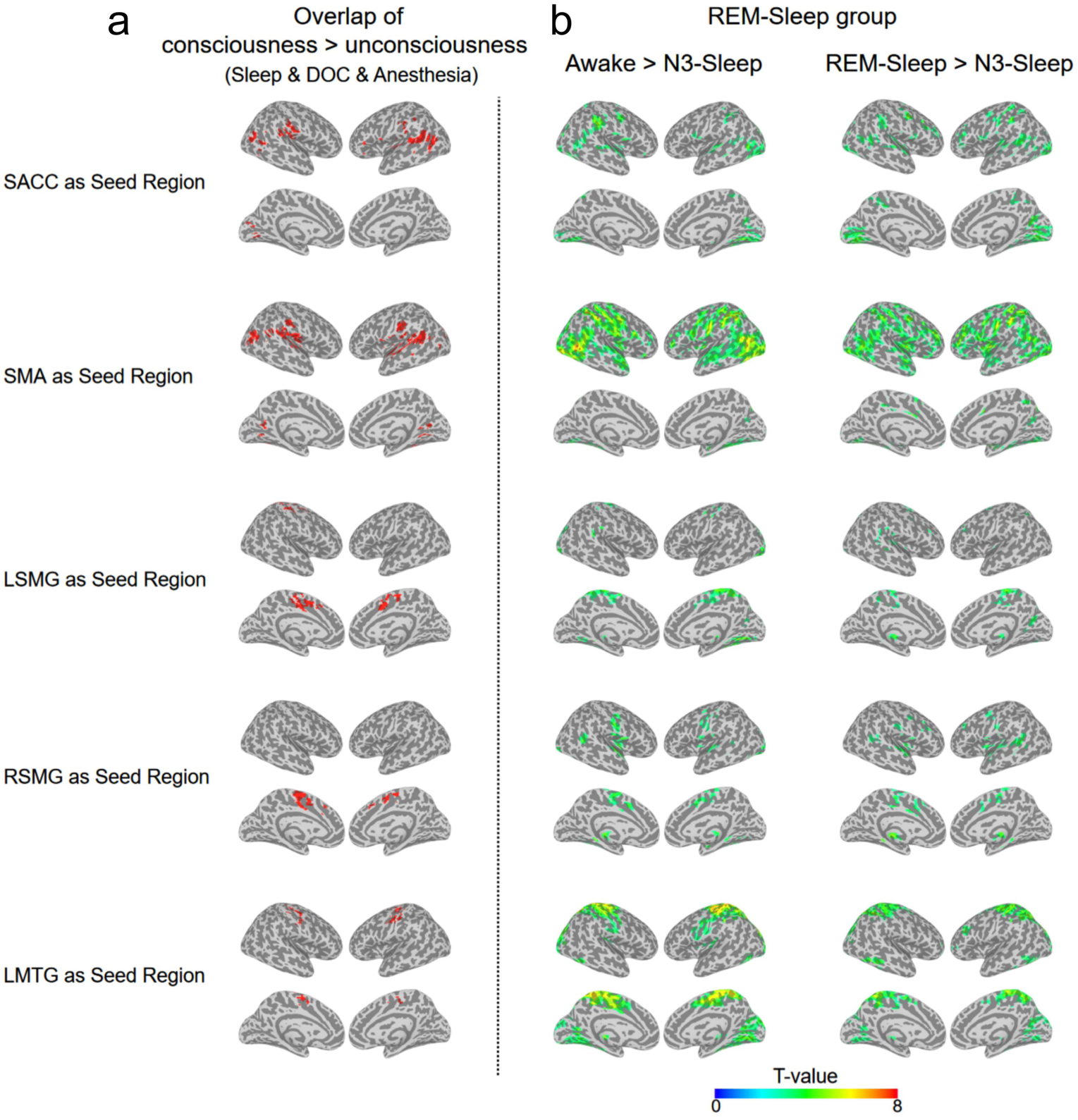
Reduced voxel-wised functional connectivity in unconscious states (a) For each seed region, the overlapped regions for all consciousness vs. unconsciousness contrasts (BL > UWS; MCS > UWS, awake > N3-sleep, N1-sleep > N3-sleep, and consciousness > anesthesia) (p < 0.005 uncorrected, volume > 20 voxels). (b) In sleep group 2, the voxel-wised FCs contrast: Awake > N3-sleep (the left panel) and REM-sleep > N3-sleep. The activation clusters are displayed (p < 0.005 uncorrected, volume > 20 voxels).

**Supplementary fig. 7.**
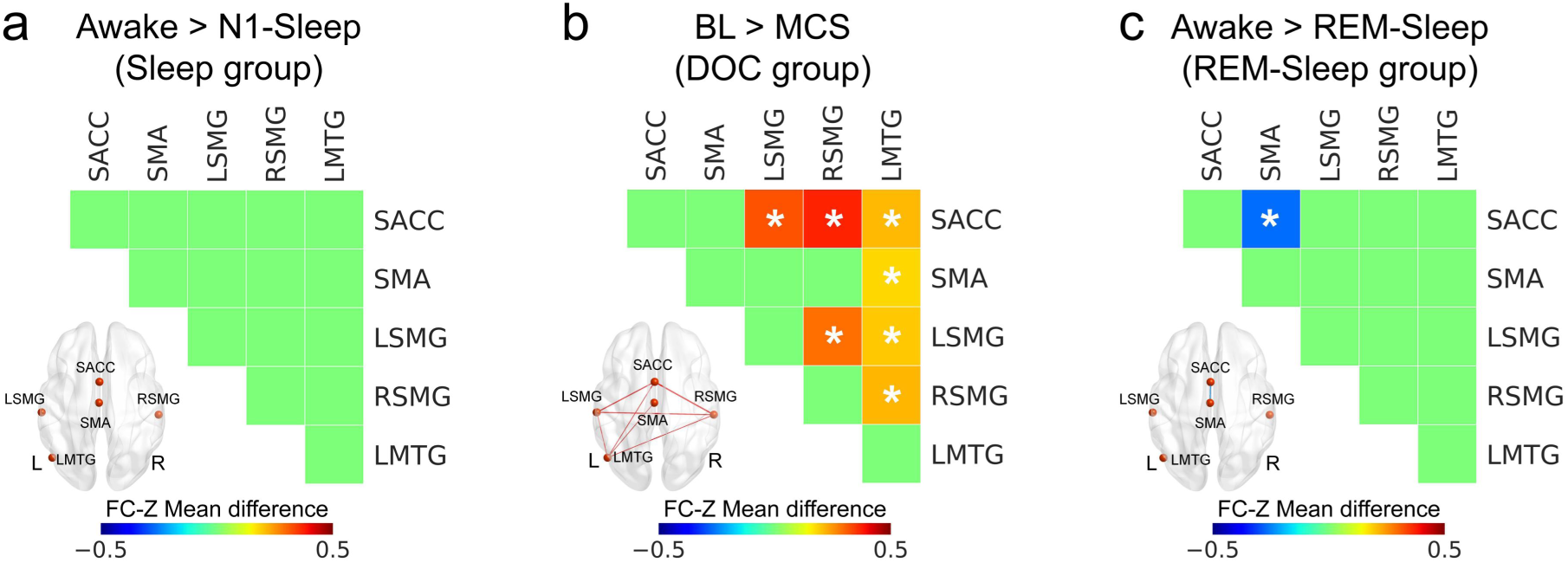
The functional connectivity difference between different conscious states. (a) No difference between awake and N1-sleep in sleep group. (b) The difference between BL and MCS in DOC group. (c) The difference between awake and REM-sleep in REM-sleep group. * means p < 0.05 FDR corrected. SACC = supragenual anterior cingulate cortex; SMA = supplementary motor area; LSMG = left supramarginal gyrus; RSMG = right supramarginal gyrus; LMTG = left middle temporal gyrus.

**Supplementary fig. 8.**
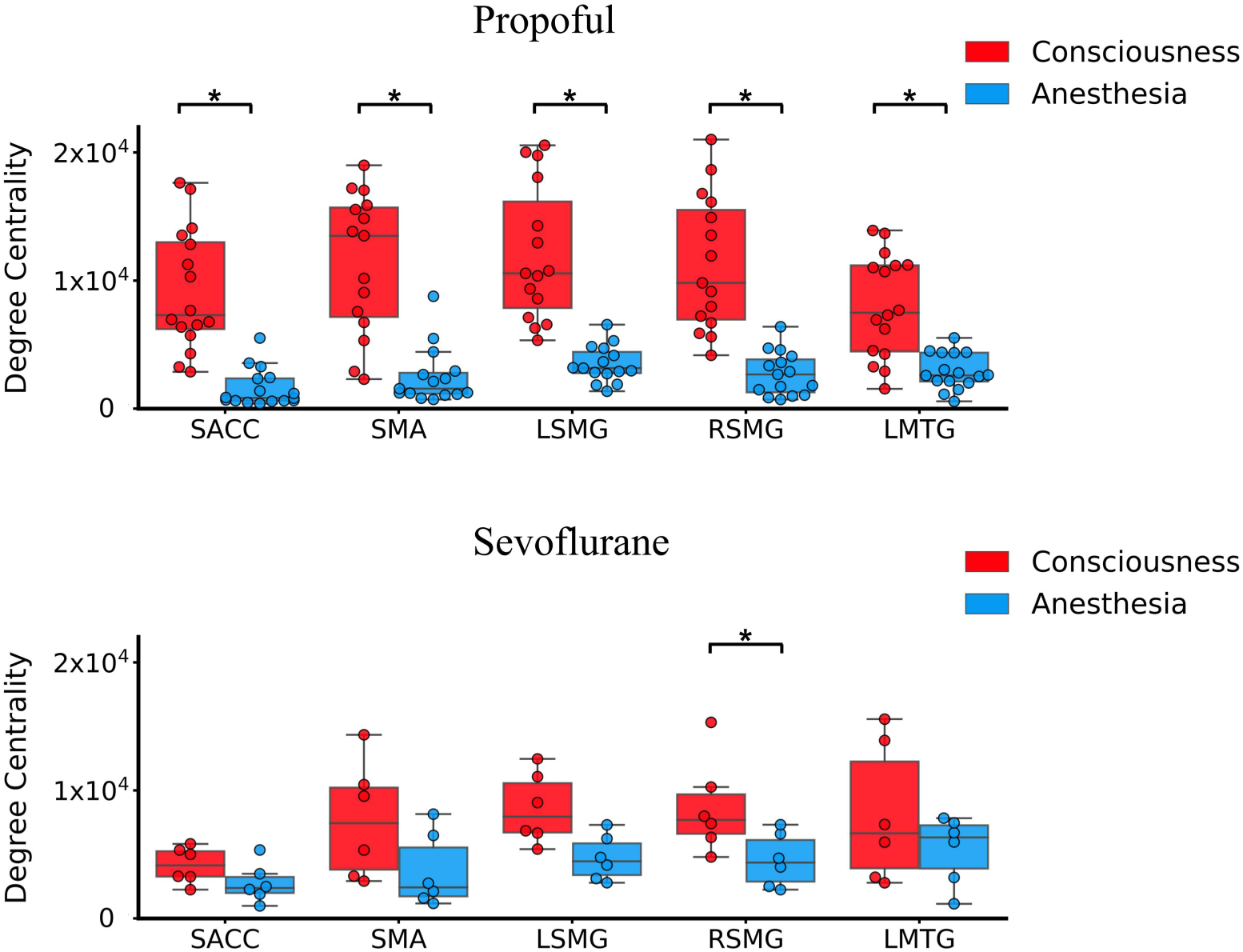
The degree centrality for participants adopting Propofol and Sevoflurane. SACC = supragenual anterior cingulate cortex; SMA = supplementary motor area; LSMG = left supramarginal gyrus; RSMG = right supramarginal gyrus; LMTG = left middle temporal gyrus. * means p < 0.05 FDR corrected.

## Notes

### Competing Interest Statement

The authors have declared no competing interest.

